# Plio-Pleistocene decline of mesic forest underpins diversification in a clade of Australian *Panesthia* cockroaches

**DOI:** 10.1101/2024.05.30.596734

**Authors:** Maxim W.D. Adams, James A. Walker, Harley A. Rose, Braxton R. Jones, Andreas Zwick, Huiming Yang, James Nicholls, Diana Hartley, Stephen Bent, Nicholas Carlile, Ian Hutton, Simon Y.W. Ho, Nathan Lo

**Author notes:** **Correspondence** Maxim Adams, LEES Building F22, School of Life and Environmental Sciences, University of Sydney, Sydney NSW 2006, Australia Nathan Lo, LEES Building F22, School of Life and Environmental Sciences, University of Sydney, Sydney NSW 2006, Australia.

## Abstract

The progressive aridification of the Australian continent, and coincident decline of mesic forest, has been a powerful driver of allopatric and environmental speciation in native species. The relictual mesic forests of the eastern seaboard now harbor a diverse group of endemic fauna, including the wood-feeding cockroaches of the genus *Panesthia*, which reached the continent via two separate invasions from Melanesia. The more recent of these colonization events gave rise to a group of five recognized species, occurring in mainland woodlands, sclerophylls and rainforests, as well as the forests and grasslands of the Lord Howe Island Group. Due to limited sampling in molecular studies and doubt regarding the standing taxonomy, there is little certainty about relationships among the species and poor understanding of the effects of ancient climatic changes upon their evolution. We undertook a comprehensive phylogenetic analysis of the clade, using complete mitogenomes and nuclear ribosomal markers from nearly all known morphospecies and populations. Our time-calibrated phylogenetic analyses reveal six unrecognized, highly divergent lineages, and suggest that these have arisen primarily through vicariance as rainforests fragmented during Plio-Pleistocene glacial cycles (2–5 million years ago). Ancestral niche reconstructions also evidence a tropical rainforest origin for the group, followed by at least three niche transitions into drier forest, including one associated with the singular colonization of the Lord Howe Island Group. Finally, we find evidence of frequent, parallel wing reduction, in potential association with the contraction of forest habitats into small refugia. Our results reiterate the far-reaching role of ancient aridification in driving speciation, niche expansion and morphological evolution in Australian fauna.

## 1. Introduction

The mesic forests of eastern Australia are among the most biodiverse habitats in the world (Ebach, 2017; Williams et al., 2011b). Occurring in a peri-coastal distribution from Queensland to Victoria, these forests are the fragmented relics of the Gondwanan rainforests that once spanned the entire continent (Hill, 1994; White, 1986, 1994). This fragmentation, caused by the gradual aridification of the Miocene and the dramatic arid cycles of the Plio-Pleistocene, saw rainforests progressively replaced by drier sclerophyll and woodland elements, with transformative effects upon the mesic biota (Bryant and Krosch, 2016; Byrne et al., 2011; Harvey et al., 2017). Untangling the complex dynamics of vicariance, adaptation and niche transition remains a central goal in Australian biogeography.

A group of consequent interest are the saproxylic (dead wood-feeding) cockroaches of the genus *Panesthia*. Originating in Asia, the *Panesthia* invaded Australia in two independent waves in the middle and late Miocene, following the collision of the Sahul and Sundaland tectonic plates (BeasleyLHall et al., 2021b; Lo et al., 2016; Maekawa et al., 2003). The latter of these colonizations has attained a broad geographic range spanning mesic, sclerophyll and woodland forests across the Australian eastern seaboard, as well as the Lord Howe Island Group (LHIG), a volcanic archipelago ∼600 km east of New South Wales (Beccaloni, 2014; Roth, 1977). Intriguingly, the insects themselves are highly sedentary and typically reside long-term within decaying logs. Many lineages have also undergone degrees of secondary wing reduction, rendering them completely flightless; while full-winged individuals manually remove their wings soon after the final molt, thereby losing flight capacity (Bell et al., 2007b; O’Neill et al., 1987). Their low vagility and habitat specificity position the group (hereafter “*Panesthia*”) as sensitive biogeographic indicators, and raise the question of how they were impacted by the fragmentation of mesic forest.

The clade comprises an ‘archipelagic’ array of isolated mesic lineages; as well as less wet-adapted species that have broader, contiguous distributions through the surrounding matrix of drier sclerophyll or open woodlands. The origins of the mesic populations remain contentious, with two broad models advanced across the literature. Based on the present habitats of Melanesian relatives, it has been suggested that the ancestral *Panesthia* were rainforest obligates (Maekawa et al., 2003), which presumably diversified through vicariance as mesic environments fragmented. This explanation would be consistent with many paradigmatic examples of allopatric speciation across the mesic biome (e.g. Bell et al., 2007a; Moreau et al., 2015; Oberski et al., 2018; Rix and Harvey, 2012). However, the distribution of mesic *Panesthia*, spanning the entire eastern seaboard, is unusually broad for a dispersal-limited invertebrate (reviewed by Bryant and Krosch, 2016), and their ancestral habitat remains poorly characterized. An alternative explanation is that the ancestors of the extant species had already come to inhabit dry sclerophyll or woodland, and subsequently colonized individual rainforest refugia as they dispersed (BeasleyLHall et al., 2021b). Transitions from dry, even arid, habitat to rainforest have been reported in a wide range of species (Byrne et al., 2018), including sedentary invertebrates (e.g., Rix et al., 2021). Interestingly, the most widespread taxon within the group, *Panesthia cribrata*, is known from both wet and dry sclerophyll, and shows a close morphological affinity to multiple, geographically separated rainforest lineages.

The most divergent habitat niche is now observed in *P. lata*, which is endemic to the LHIG. Estimated to have colonized the archipelago 2–6 million years ago (Lo et al., 2016), the species occupies rainforest on Lord Howe Island itself, and more exposed grasslands on several of the surrounding islets. The cockroaches are also unique in constructing shallow burrows under stones or leaf litter, rather than inhabiting logs (Rose, 2003). It has been suggested that the offshore islet populations expanded their niche *in situ* in response to the exposed conditions of the archipelago (BeasleyLHall et al., 2021b). However, without a clear understanding of the ancestral habitat of mainland congeners, and of the species itself, the evolution of environmental tolerances of *P. lata* remains opaque.

To date, biogeographic understanding has been limited by systematic irresolution. The monophyly of the *Panesthia* is well supported, yet estimates of the relationships within the group have been highly unstable between studies (BeasleyLHall et al., 2021b; Legendre et al., 2017; Legendre et al., 2015; Lo et al., 2016). While five species are presently recognized, there is mounting evidence of discordance between genetic results and the morphology-based taxonomy, suggesting cryptic speciation and phenotypic parallelism (BeasleyLHall et al., 2021b; Djernæs et al., 2020; Lo et al., 2016). Previous investigations have included only few representatives of the clade and have lacked the resolution to confidently delimit species or interrogate geographic patterns of diversity.

Here we undertake the first comprehensive phylogenetic investigation of the *Panesthia*, including novel populations never previously examined in taxonomic or genetic studies. In clarifying their systematics and evolutionary history, we explore three fundamental questions: 1) Did the contemporary distribution of mesic lineages arise through vicariance, or were these habitats separately colonized by dry-forest ancestors?; 2) Were the ancestors of *P. lata* pre-adapted to the drier conditions of the LHIG, or did the species expand its niche following island colonization?; and 3) Do flightless morphs share a common ancestry, or has wing reduction occurred in parallel? The results of our study highlight the complex evolutionary dynamics associated with Australia’s mesic biome.

## 2. Methods

### 2.1. Sampling and DNA sequence data

Samples of mainland taxa were collected between 1998 and 2023 from across New South Wales and Queensland, Australia. Specimens were stored in 70–100% ethanol or in pinned collection prior to DNA extraction, and are presently held in the private collections of H.A. Rose and J.A. Walker. Due to the scarcity of *P. lata* in collection, we retrieved specimens from the LHIG in July and August of 2022, from Blackburn Island, Roach Island and a newly discovered relict population on Lord Howe Island (Adams et al., in prep.). In supplement, we subsampled tissue from historical specimens of *P. lata* collected on Lord Howe Island and Ball’s Pyramid between 1869 and 1973, sourced from the Australian Museum, Sydney and the Macleay Museum, Sydney. A full list of material with GenBank accession numbers is provided in Supplementary Table S1.

DNA was extracted from leg muscle tissue to avoid contamination from *Blattabacterium* bacterial endosymbionts, which occur in abdominal fat bodies (Kinjo et al., 2015). DNA sequencing was outsourced to the Australian National Insect Collection, Canberra, utilizing an approach suitable for highly fragmented historical DNA (see Jin et al., 2020; Zwick and Zwick, 2023 for methods). In summary, genomic DNA was extracted using proteinase K digestion and a silica filter-based approach in a 384-well format. Up to 5 ng of extracted DNA were used to build ligation-based whole-genome shotgun DNA sequencing libraries, utilizing an acoustic liquid handler (Echo 525; Beckman Coulter, California, USA) to miniaturize reaction volumes for increased reaction efficiency. DNA libraries of different samples were pooled equimolar and sequenced at the Australian National University’s Biomolecular Resource Facility on an Illumina NovaSeq 6000 platform, using an S1 flow cell and a 300-cycle sequencing kit. The output of 1.65 billion 150 bp paired-end reads were demultiplexed to yield a median of 27,321 reads per sample. Output data included reads from most or all of the mitochondrial genome (hereafter “mitogenome”), as well as the nuclear ribosomal operon (which encodes 18S, 5.8S and 28S ribosomal RNA, alongside the internal transcribed spacers *ITS1* and *ITS2*).

We assembled raw reads into contigs using SPAdes v.3.12.0 (Bankevich et al., 2012) with default settings and sampling *k* values of 33, 55, 77, 91, and 121. To generate mitogenomes, contigs produced in SPAdes were imported into Geneious Prime v.2022.1.1 (https://www.geneious.com) and assembled to a reference sequence using the Map to Reference tool with default settings and medium sensitivity. Where available, we extracted the single contig comprising the near-complete mitogenome (*ca.* 15,000 bp). Otherwise, we extracted the consensus sequence of multiple contigs, selecting the bases with the highest representation. Reference sequences were chosen to represent the closest known sister taxon to each sample, based on the phylogenetic framework of Beasley-Hall et al. (2021b); these were sourced from GenBank or from mitogenomes generated presently.

We annotated mitogenomes using the MITOS web server (Bernt et al., 2013) under default settings for invertebrate mitochondrial DNA. Duplicated annotations and split genes were corrected in Geneious, and we checked for errors against published references for *Panesthia parva*, *Panesthia sloanei* and *Panesthia angustipennis* (BeasleyLHall et al., 2021b). We then aligned the sequences for each gene individually using the MUSCLE algorithm (Edgar, 2004) within Geneious. The control region was omitted from our analyses, as it includes repetitive DNA regions that are not reliably assembled from short reads (Bourguignon et al., 2018). Nuclear markers were generated via the same steps, initially using a nuclear ribosomal operon reference sequence for *Panesthia angustipennis* (comprising, in order, *18S*, *ITS1*, *5.8S*, *ITS2* and *28S*; Che et al., 2022).The final mitochondrial and nuclear alignments represented a total of 14,707 bp and 5,596 bp, respectively, and were analysed separately to account for potential genealogical discordance.

One novel morphospecies, known from Koombooloomba State Forest, Queensland, was discovered after the main round of sequencing. We used polymerase chain reaction (PCR) amplification to generate a 604 bp fragment of mitochondrial *CO1* and a 352 bp fragment of mitochondrial *16S* from a single specimen (see Supplementary Table S2 for primers and PCR protocols). Econotaq^TM^ master mix (New England Biolabs, Massachusetts, USA) was used as the source of free nucleotides and reaction buffers. PCR products were cleaned using Exosap-IT (Thermo Fisher Scientific, Massachusetts, USA) and sent to Macrogen (Seoul, Gyeonggi, South Korea) for Sanger sequencing.

In total, 137 ingroup taxa were sequenced successfully. We combined these sequences with a complete mitogenome of the ingroup taxon *Panesthia parva*, and six outgroup representatives of the Australian and Melanesian Panesthiinae, which were retrieved from GenBank (Supplementary Table S1). Each gene alignment was manually checked for reading frames and premature stop codons in Seqotron v.1.0.1 (Fourment and Holmes, 2016), and ambiguously aligned regions were removed. We then tested for substitutional saturation using Xia’s method in DAMBE7 (Xia, 2018), and in its absence retained all codon positions for analysis.

### 2.2. Phylogenetic analysis

For the mitochondrial data set, we opted for a biologically relevant partitioning scheme consisting of first, second and third codon positions of protein-coding genes, rRNAs and tRNAs, in accordance with previous phylogenomic studies of the Blattodea (BeasleyLHall et al., 2021b; Bourguignon et al., 2014; Bourguignon et al., 2018; Cameron et al., 2012). However, we modified the scheme by implementing a separate partition for *CO1*, to enable calibration of the molecular clock (for a total of six partitions). We used the ModelFinder function (Kalyaanamoorthy et al., 2017) to infer the best-fitting substitution model for each partition, based on Bayesian information criterion scores (Supplementary Table S3). For the nuclear data set, we co-estimated the optimal scheme and substitution model using ModelFinder (2 partitions: *18S*+*5.8S*+*28S*, *ITS1+ITS2*; Supplementary Table S3). This scheme was also modified for molecular dating by implementing a separate partition for *28S* (for a total of three partitions). Preliminary analyses showed that the segregation of *CO1* and *28S* did not affect the inferred tree topologies.

Maximum-likelihood (ML) phylogenetic analyses were performed in IQTREE v.2.2.2 (Minh et al., 2020). Node support was estimated using 10,000 ultrafast bootstrap replicates (UFBoot; Hoang et al., 2018) and the SH-like approximate likelihood-ratio test (SH-aLRT) with 1,000 iterations. Following recommendations of the package, we considered UFBoot > 0.95 and SH-aLRT > 0.8 to indicate strong support.

We undertook two additional analyses to investigate the discordance of the mitogenomic and nuclear topologies (see Results). First, we tested whether the signal from the nuclear data set was significantly inconsistent with the mitogenomic results. Using IQTREE, we analysed the nuclear markers under topology constraints following the species-level branching order of the mitogenomic ML topology, and statistically compared its adequacy relative to the unconstrained nuclear tree using six different metrics (Supplementary Table S4). Second, to assess the relative information content of the two data sets, we estimated ML and Bayesian trees using a concatenated alignment comprising all mitochondrial and nuclear loci. Partitions and substitution models were as described previously.

### 2.3. Molecular dating

Phylogenetic trees and evolutionary timescales were jointly estimated using Bayesian inference in BEAST v.10.4 (Suchard et al., 2018). We modeled among-lineage rate variation using an uncorrelated lognormal relaxed clock (Drummond et al., 2006) and specified a birth-death tree prior, which is most appropriate for the combination of interspecific and intraspecific sampling in the data set (Ritchie et al., 2016). Each partition was assigned a separate GTR+I+G substitution model, representing the closest model to those estimated in ModelFinder. We ran two independent chains, drawing samples every 10^3^ steps until convergence was observed in Tracer v.1.7.2 (Rambaut et al., 2018) and the effective sample size for each parameter reached ≥ 200 (1.5×10^8^ steps for the mitogenomic data set, 1×10^8^ steps for the nuclear data set). The maximum-clade-credibility tree was generated in TreeAnnotator with a 10% burn-in.

There are no known fossil calibrations proximal to the Mio-Pliocene divergences inferred for the *Panesthia* (BeasleyLHall et al., 2021b; Lo et al., 2016). We consequently utilized two different techniques to estimate the evolutionary timescale. First, we followed BeasleyLHall et al. (2021a) to specify an informative prior distribution for the substitution rate, based on the late Miocene diversification of Mediterranean *Dolichopoda* crickets (Allegrucci et al., 2011). The genus is ecologically and biologically similar to the *Panesthia*, comprising nonvagile, subterranean species of comparable body size (Allegrucci et al., 2021). In addition, the diversification of *Dolichopoda*, initiating *ca.* 7 Ma, is temporally proximal to the Mio-Pliocene radiation of the *Panesthia* estimated by BeasleyLHall et al. (2021b). For the mitogenomic analysis, we specified a substitution rate prior for *CO1* (1.6×10^-2^ ± 1.1×10^-4^ substitutions/site/Myr), due to its conserved substitution rate across insect orders (Gaunt and Miles, 2002; Papadopoulou et al., 2010). For the nuclear analysis, the only available rate estimate was for *28S* (6.4×10^-4^ ± 4×10^-6^ substitutions/site/Myr). These were applied as normal priors with uncertainty corresponding to the standard deviation, and the rates of remaining partitions were estimated during analysis.

For our second approach, we applied two secondary calibrations to the backbone nodes of the tree, based on the evolutionary timescale inferred by BeasleyLHall et al. (2021b). Using the mitogenomic data set, we specified normal priors for the age of the root (29.18 ± 4.06 Ma) and for the node uniting *Panesthia* with the *P. angustipennis* complex (25.02 ± 5.42 Ma). Because the date estimates in BeasleyLHall et al. (2021b) were inferred using ancient fossil calibrations, which tend to artifactually deepen shallow nodes (Hipsley and Müller, 2014; Ho et al., 2011), we consider the second approach to represent a conservative upper bound and primarily focus on results from the *Dolichopoda* rate of evolution.

### 2.4. Species delimitation

To investigate the presence of unrecognized species diversity, we followed a two-step process to delimit highly divergent, monophyletic clades (hereafter “operational taxonomic units”, or OTUs; see Supplementary Material for full methods). First, we generated OTU hypotheses *de novo* from the mitogenomic data set using the Generalized Mixed Yule Coalescent (GMYC; Zhang et al., 2013) and the distance-based Assemble Species by Automatic Partitioning (ASAP; Puillandre et al., 2021). Then, in accordance with best practice outlined by Carstens et al. (2013), we validated the output delimitations in Bayesian Phylogenetics and Phylogeography v.4.1.4 (BPP; Yang, 2015), using both the mitogenomic and nuclear data sets.

### 2.5. Historical biogeography and niche evolution

We reconstructed ancestral geographic ranges and habitat niches using the ultrametric phylogeny inferred in BEAST, with the input tree pruned to include a single representative of each OTU. Due to the instability of the nuclear topology across analyses (see Results), reconstructions were only performed with the mitochondrial data set.

We explored changes in geographic range with the R package BioGeoBEARS (Matzke, 2013). Taxa were divided among five biogeographic zones: North Queensland, Central Queensland, Mid-Eastern Australia, South-Eastern Australia, and the LHIG. These correspond to zoogeographical subregions outlined by Ebach et al. (2013), which reflect well-documented partitions in the distributions of eastern seaboard fauna. Colonization of the LHIG was assumed to be unidirectional, while all other transitions were coded as equiprobable and time-constant. Based on contemporary ranges, the maximum number of areas was set as two.

We compared three widely applied biogeographic models, each with unique assumptions about the cladogenetic and anagenetic processes underpinning speciation. These were Dispersal-Extinction-Cladogenesis (DEC), Dispersal-Vicariance analysis (DIVA-like) and Bayesian Analysis of Biogeography (BayArea-like). Models were run under default parameters and compared using the corrected Akaike information criterion (AICc). We opted not to include the jump dispersal parameter (+J) due to ongoing debate regarding its statistical validity (Matzke, 2022; Ree and Sanmartín, 2018).

We also explored the timing of transitions between wet, dry, and open forest using ancestral niche reconstruction in the R package *Nichevol* v.1.19 (Owens et al., 2020). Species were split across three major habitat categories (following Braby et al., 2020; Mitchell et al., 2014): closed forest (rainforest, wet sclerophyll or vine thicket; canopy cover > 80%, precipitation:evaporation > 0.4), open forest (dry sclerophyll and scrub; canopy cover 50– 80%) and woodland (canopy cover < 50%). We then modeled the change in habitat along the phylogeny using a maximum-likelihood framework. To avoid overestimating niche lability (Barve et al., 2011; Owens et al., 2020), *Nichevol* allows the presence of a taxon in a niche to be coded as “uncertain”. We considered a niche to be uncertain unless the species was confirmed to be absent from the niche in an area adjacent to its known habitat. Ecological data were compiled from the personal observations of H. A. Rose and J. A. Walker and validated against the Cockroach Species File Online database (Beccaloni, 2014).

### 2.6. Evolution of flight loss

Lastly, we modeled the evolution of wing morphology using ancestral state reconstructions (ASRs). We classified OTUs as fully winged (macropterous), partially winged (brachypterous) or vestigially winged (micropterous) (see Figure 4 for representative images). Present-day characters were mapped onto our mitogenomic species tree, and ASRs were performed in the R package *phytools* v.2.1.1 (Revell, 2012). We reconstructed changes along the topology using a continuous-time Markov chain model, and estimated the most likely state at each node. The *Panesthia* are understood to have originated from macropterous ancestors (Lo et al., 2016), thus we included the closest known outgroup species *P. angustipennis angustipennis*, and fixed the root of the tree to be macropterous. We undertook two reconstructions, the first with a unidirectional rate matrix permitting only evolution away from the ancestral state (since wing re-evolution is considered highly unlikely; Trueman et al., 2004). Our second reconstruction permitted free transitions between all character states (by specifying a bidirectional “all rates different” model). As one OTU was found to be highly polymorphic, we undertook separate analyses reconstructing wing evolution within the OTU, using a single representative from each sampling locality.

## 3. Results

### 3.1. Phylogenetic relationships and species delimitation

Our analyses uniformly support the monophyly of the *Panesthia*. Maximum-likelihood and Bayesian analyses of the mitogenomic data set produced near-identical estimates of the phylogeny, differing only in the placements of some terminal branches (Figure 1). Genetic relationships within and between major lineages were generally clustered by geographic locality, with the northern Queensland samples (Clades A+B + *P.* sp. Koombooloomba) forming successive sister groups to a clade comprising samples from central and southern Queensland, New South Wales, and the LHIG (Clade C).

**Figure 1.**
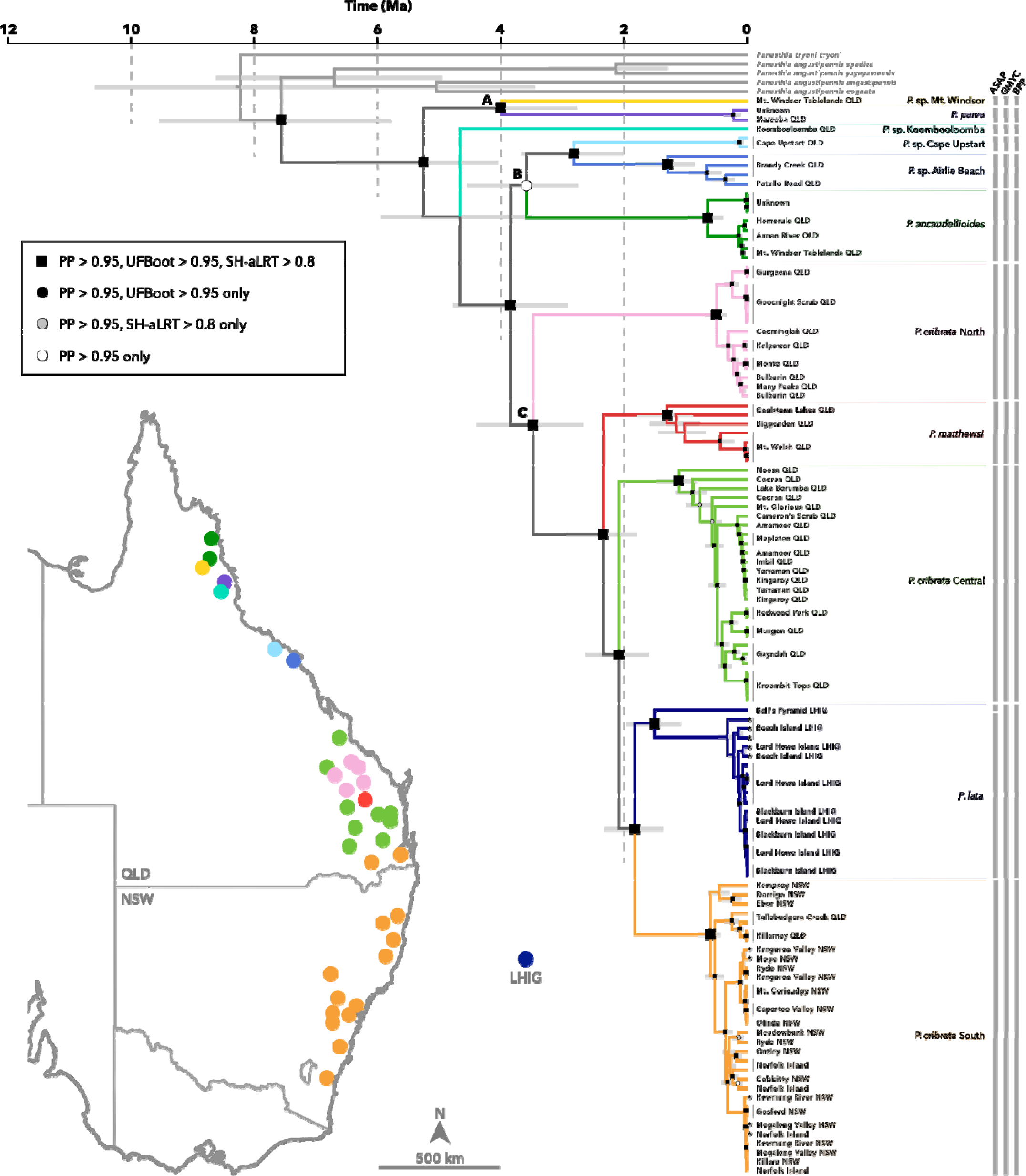
Dated phylogeny of the *Panesthia* inferred from complete mitochondrial genomes in BEAST and IQTREE. The evolutionary timescale was inferred using a previous estimate of the *CO1* substitution rate (Allegrucci et al. 2011). PP: posterior probability, UFBoot: ultrafast bootstrap, SH-aLRT: SH-like approximate likelihood ratio test, QLD: Queensland, NSW: New South Wales, LHIG: Lord Howe Island Group. Size of node labels varied for visual clarity. Letters A–C denote crown nodes of major clades. Stars denote tips with varying position between BEAST and IQTREE analyses. Results of species delimitation analyses are displayed to the right of tips. GMYC and ASAP results are based on mitochondrial genomes only, while BPP results are based on mitochondrial genomes and the nuclear ribosomal operon. Nomenclature and coloration of operational taxonomic units follows BPP results. **Inset:** distribution of sampling localities in eastern Australia.

The three species delimitation methods applied to this data set yielded similar estimates of OTUs: 10 from GMYC, 12 from ASAP and 11 from BPP (Figure 1). Since the BPP analysis incorporated both mitochondrial and nuclear markers, and produced delimitations that best reflect accepted species boundaries, we consider the 11-OTU scheme to be the most robust and refer to this in subsequent sections.

Four of the OTUs correspond to novel localities previously unsampled in taxonomic or genetic studies: *P*. sp. Cape Upstart, *P.* sp. Airlie Beach, *P*. sp. Mt. Windsor and *P.* sp. Koombooloomba. Further, *P. cribrata* was found to comprise three divergent clades, which we presently refer to as *P. cribata* North, *P. cribrata* Central and *P. cribrata* South. The monophyly of each OTU was well supported (excluding *P.* sp. Koombooloomba and *P.* sp. Mt. Windsor, each of which was represented by a single sample). The phylogenetic positions of all OTUs were also resolved with uniformly high support (PP, UFBoot, SH-aLRT > 0.95), excepting the crown node of Clade B (PP = 0.97, UFBoot = 0.9, SH-aLRT = 69.5) and the node uniting *P.* sp. Koombooloomba with its sister group (PP = 0.91, UFBoot = 0.97, SH-aLRT = 84.3; but note that only two markers were sequenced for this sample).

The mitochondrial OTUs were consistently recovered as monophyletic in our analyses of the nuclear ribosomal loci, with the exception of *P. cribrata* Central and *P. cribrata* South, which were paraphyletic in the ML and Bayesian trees, respectively (Figure 2, Supplementary Figure S1; note that *P.* sp. Koombooloomba was absent from the nuclear data set). However, the branching order of the OTUs and the populations within OTUs were somewhat discordant with the mitochondrial results, and differed between ML and Bayesian analyses (in both cases with relatively low node support; Figure 2, Supplementary Figure S1). The most significant conflict was the position of *P. lata*, which was placed substantially deeper in the phylogeny, either on its own branch in ML analysis (UFBoot = 46, SH-aLRT = 60.7) or as sister to *P.* sp. Airlie Beach in Bayesian analysis (PP = 0.95).

**Figure 2.**
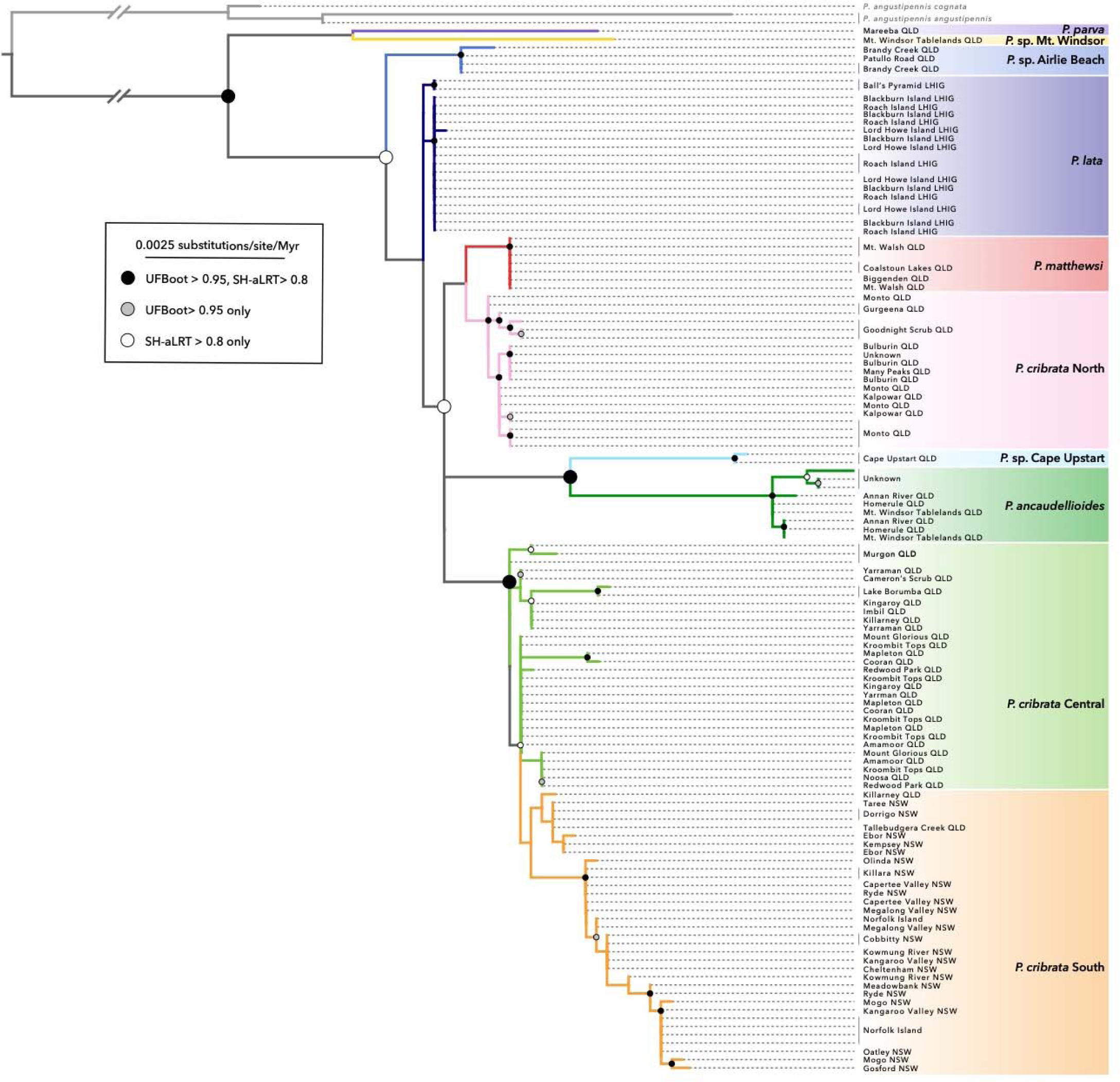
Maximum-likelihood phylogeny of the *Panesthia* inferred from the nuclear ribosomal operon in IQTREE. UFBoot: ultrafast bootstrap, SH-aLRT: SH-like approximate likelihood-ratio test. Sizes of node labels varied for visual clarity. Nomenclature and coloration of operational taxonomic units follows the mitogenomic (BPP) results.

The constrained nuclear tree, which followed the mitogenomic branching order, was strongly rejected by all six metrics in IQTREE in favor of the unconstrained tree (Supplementary Table S4), indicating that the phylogenetic signal from the nuclear data is significantly incongruent with the mitogenomic topology. However, when we estimated an ML tree from a concatenated alignment comprising all available loci, the species-level topology was almost identical to the mitogenomic tree, differing solely in the placement of *P. lata* as sister to *P. cribrata* Central + South (UFBoot = 98, SH-aLRT = 99.8; Supplementary Figure S2).

### 3.2. Molecular dating

Based on *CO1* rate calibration of the mitogenomic data set, the stem age of the *Panesthia* was placed at *ca.* 7.62 Ma (95% credible interval [CI] 5.79–9.53 Ma), with a crown age of *ca.* 5.30 Ma (95% CI 4.06–6.66 Ma). The stem ages of all eleven OTUs were dated to the Pliocene or early Pleistocene (*ca.* 1.8–5 Ma), with intraspecific divergences occurring in the middle to late Pleistocene (< 1.5 Ma, excluding the Ball’s Pyramid lineage of *P. lata*, which diverged from conspecifics *ca.* 1.51 Ma, 95% CI 1.08–1.96 Ma).

The evolutionary timescale recovered from nuclear loci yielded a curiously deep stem age, albeit with a broad 95% CI (14.61 Ma, 95% CI 5.16–24.82 Ma). The remaining nodes had younger and less precise age estimates than in the mitogenomic analysis (Supplementary Figure S1). The crown age of the *Panesthia* was estimated as *ca.* 4.84 Ma (95% CI 2.47–7.84 Ma), while all species-level cladogenesis was estimated to have occurred in the late Pliocene and early–middle Pleistocene (*ca.* 0.5–3 Ma).

As anticipated, secondary calibration using node ages from BeasleyLHall et al. (2021b) produced a substantially deeper estimated timescale, with a stem age of 22.41 Ma (95% CI 15.46–28.89 Ma; Supplementary Figure S3). All interspecific divergences were estimated to have occurred in the middle to late Miocene (*ca.* 5.3–15.3 Ma), followed by intraspecific diversification throughout the Pliocene and Pleistocene (< 5.0 Ma).

### 3.3. Historical biogeography

In BioGeoBEARS analysis, the best-fitting model was DIVA-like (AICc = 49.64), compared with DEC (AICc = 50.78) and BayArea-like (AICc = 57.65). The ancestral state was well resolved (most probable state > 50%) for nearly all internal nodes, excluding only the clade comprising *P. lata* + *P. cribrata* South. Under this scenario, the most recent common ancestor (MRCA) of the *Panesthia* occurred in North Queensland (Figure 3). All ancestral and contemporary ranges in Clades A+B were placed in Central Queensland, while the crown node of Clade C was estimated to occur in Central Queensland alone, suggesting a southward range expansion. Presently, all mainland OTUs except *P. cribrata* South occur in Central or North Queensland, while the latter is found in Mid-eastern Australia and South-eastern Australia. The ancestral range of *P. lata*, which is presently endemic to the LHIG, could not be confidently resolved between Central Queensland, Mid-eastern Australia, or South-eastern Australia.

**Figure 3.**
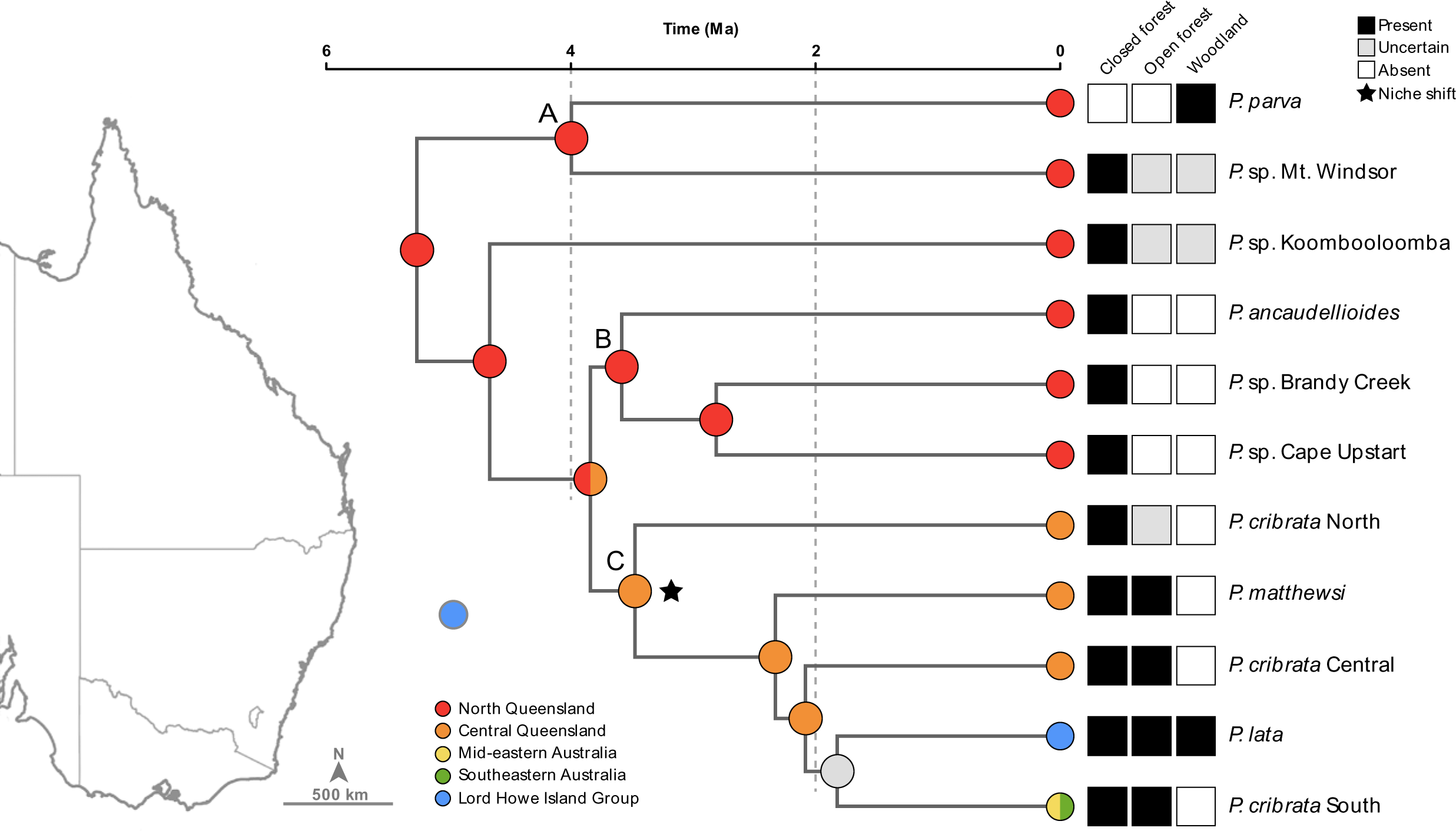
Ancestral geographic ranges and habitat niches of the *Panesthia*, reconstructed over the Bayesian chronogram inferred from whole mitogenomes in BEAST. Circles at nodes represent the most likely ancestral range, estimated using a dispersal-vicariance model in BioGeoBEARS. Multicolored circles denote a range spanning multiple bioregions, while a gray circle indicates an unresolved ancestral range. Tips are labeled with present-day distributions and habitat niches. The ancestral niche was inferred to be closed forest in *Nichevol* (not labeled), which was retained at all internal nodes until the point denoted by a star, where there was an inferred niche expansion into open forest.

Ancestral niche reconstruction in *Nichevol* suggested that the MRCA of the *Panesthia* inhabited exclusively closed forest (Figure 3). This niche was retained in the ancestors of all lineages in Clades A + B; however, we inferred a niche expansion into open forest within Clade C, occurring in the ancestor of *P. matthewsi + P. cribrata* Central + *P. lata* + *P. cribrata* South. In addition, two contemporary OTUs were found to have independently expanded or shifted their niche into woodland: *P. parva*, which inhabits woodlands only; and *P. lata*, which inhabits mesic closed forest, open forest, woodland and xeric grasslands (not labeled). No mesic lineages were found to have evolved from dry forest or woodland ancestors.

### 3.4. Wing morphology

Ancestral state reconstructions consistently found that wing reduction has occurred multiple times within the *Panesthia*. When wing re-evolution was prohibited, wing reduction was inferred to have occurred six times in total, including twice within *P.* cribrata Central (Figure 4). Five OTUs were exclusively micropterous, arising from four independent wing reductions: *P. lata*, *P. matthewsi*, *P. cribrata* North, and *P.* sp. Cape Upstart + *P.* sp. Brandy Creek (Figure 4a). In contrast, *P. cribrata* Central was found to be wing polymorphic. When wing evolution was modeled within the species, we inferred two populations to have independently become brachypterous (Murgon and Kroombit Tops) and a single population to have become micropterous (Gayndah), having evolved from a shared, brachypterous ancestor with the Kroombit Tops population (Figure 4b).

**Figure 4.**
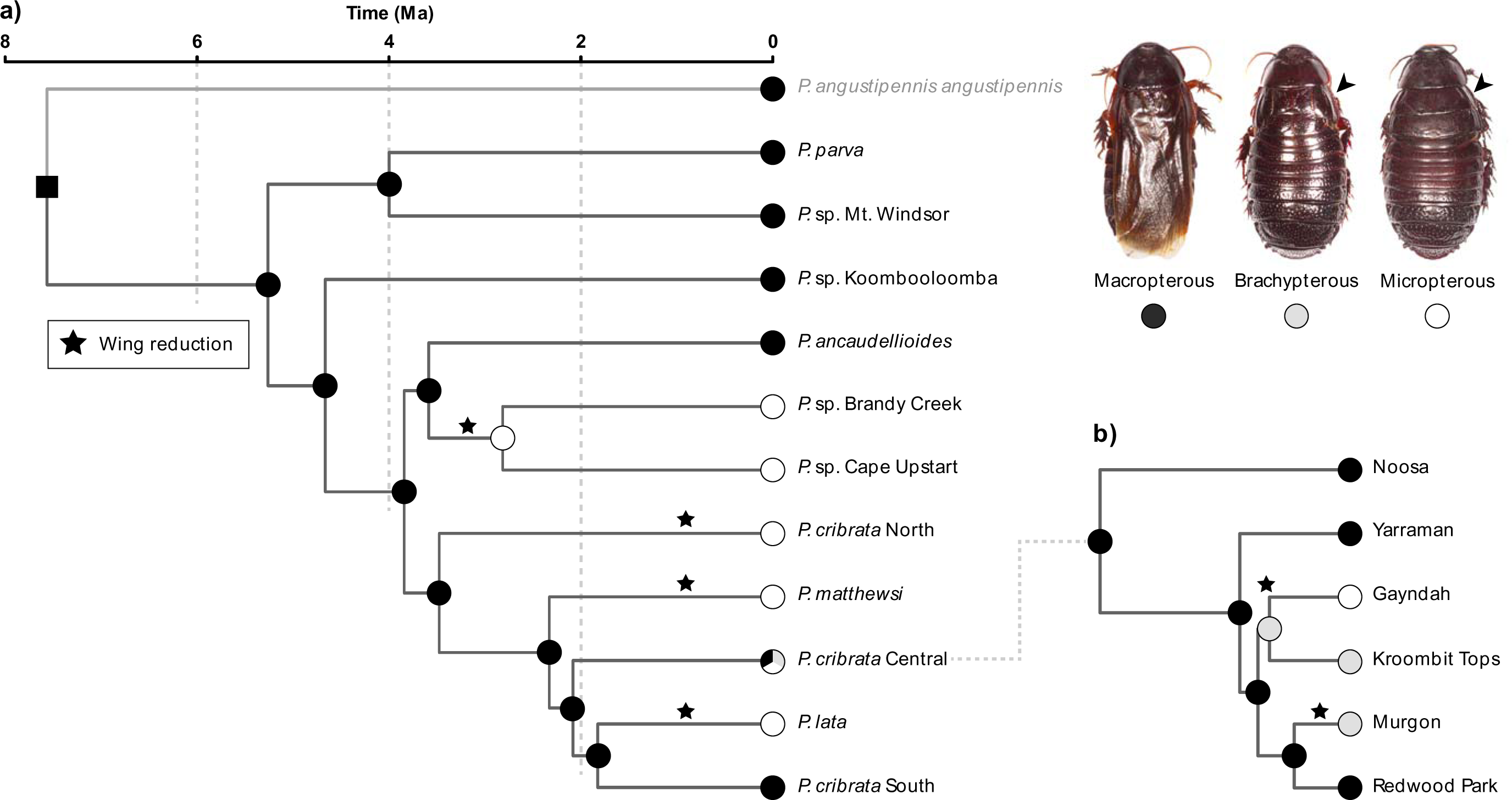
Evolution of wing morphology with wing re-evolution prohibited, reconstructed in *phytools* over the Bayesian chronogram inferred from complete mitogenomes in BEAST. **a)** Ancestral wing morphology of the 11 operational taxonomic units (OTUs). **b)** Ancestral wing morphology within the polymorphic OTU *Panesthia cribrata* Central, pruned to include a representative selection of sampling localities. Circles at nodes indicate the most probable ancestral state and the square at the root node the fixed ancestral state. States at all internal nodes were estimated with probability > 50%. Stars denote an inferred reduction of wings from a macropterous ancestor. **Inset:** representative habitus images of wing morphs. Arrows indicate reduced forewings. Photographs by Braxton Jones.

The scenario varied slightly when wing re-evolution was permitted, with an estimate of eight independent wing reductions (Supplementary Figure S4). All flightless lineages were inferred to have independently arisen from macropterous ancestors, including *P.* sp. Cape Upstart and *P.* sp. Airlie Beach, and the Gayndah and Kroombit Tops populations of *P. cribrata* Central.

## 4. Discussion

### 4.1. Phylogenetic relationships

This study substantially clarifies the evolution and systematics of the *Panesthia*. Previously thought to comprise five species, our results reveal unrecognized diversity in rainforest isolates across Queensland, and we provisionally identify eleven divergent lineages. With the notable exception of *P. cribrata*, there was strong support for the monophyly of established species in both nuclear and mitochondrial analyses. In contrast, geographic sampling was sufficient to discern three divergent mitochondrial clades within *P. cribrata*, which were polyphyletic with respect to *P. lata* and *P. matthewsi*. This potentially explains the inconsistent relationships between these species across previous studies (BeasleyLHall et al., 2021b; Lo et al., 2016), and echoes suggestions that *P. cribrata* represents a species complex (BeasleyLHall et al., 2021b; Roth, 1977). However, we are currently unable to discern morphological boundaries between the three *cribrata* lineages, and note that the non-monophyly *P. cribrata* South and Central in nuclear analyses may indicate ongoing gene flow at the boundary of their ranges. Taxonomic study is underway to determine whether and how to partition the putative complex, as well as to formally describe the novel OTUs identified by our study.

Mitogenomic analyses supported a southward grade of diversification, with all north Queensland samples forming successive sister lineages to those present in south-east Queensland and New South Wales. This is consistent with the understanding that the *Panesthia* dispersed southwards across Australia following their arrival from Melanesia (Maekawa et al., 2003). A similar overall pattern was found by Lo et al. (2016) and Beasley-Hall et al. (2021b), using primarily mitochondrial markers with lower taxon sampling. Although the nuclear topologies also supported a north–south geographical gradient, the trees were less strictly geographically clustered, and estimated an early divergence of *P. lata*. This aligns with the results of Legendre et al. (2015, 2017), where *P. lata* was inferred to be sister to a clade uniting *P. cribrata* and *P. ancaudellioides*, based on analysis of concatenated mitochondrial and nuclear genes.

Mito-nuclear discordance is not unusual in data sets spanning the boundary between species- and population-level processes, and could indicate incomplete lineage sorting, selection or introgression. Nuclear introgression would be consistent with the male-biased dispersal observed in *Panesthia* species (O’Neill et al., 1987). However, given the slower evolution and low node support of the nuclear phylogenies, it is likely that much of the discordance is due to low phylogenetic information content (Zink and Barrowclough, 2008). This interpretation is broadly supported by our analyses of the concatenated data set, which produced a topology almost identical to that inferred from mitogenomes. Therefore, we focus our discussions on the mitogenomic results, noting well-supported discrepancies where relevant. In future, the extent of introgression could be investigated using a wider suite of nuclear markers, such as single-nucleotide polymorphisms.

### 4.2. Mainland biogeography

The *Panesthia* were found to have diverged from Melanesian ancestors in the late Miocene (*ca.* 7.62 Ma). By this point, paleobotanic and phylogeographic data suggest that rainforest had substantially retracted, with both mesic and sclerophyllous elements occurring across the eastern seaboard (Byrne et al., 2011; Martin, 2006; Rix and Harvey, 2012). Nonetheless, the ancestral *Panesthia* were inferred to be wet-forest obligates. A rainforest origin is concordant with the ecology of the genus in Southeast Asia (Roth, 1979b; Wang et al., 2014), although the precise relationships between Australian and Melanesian species are unclear because much of the latter’s diversity remains undescribed. A wider body of evidence also indicates that faunal exchange from Asia was dominated by rainforest species (e.g., Braby et al., 2020; Rowe et al., 2011; Roycroft et al., 2022), suggesting that paleobotanic conditions were favorable to their dispersal.

The *Panesthia* now attain their greatest diversity in Queensland, comprising nine potential species. In sharp conflict with the hypothesis that rainforest patches were colonized by woodland-adapted ancestors (BeasleyLHall et al., 2021b; Roth, 1977), eight of these were found to have either retained the ancestral closed forest niche or only expanded into open forest (i.e., broadly remained within the mesic biome), with only a single transition to open woodland in the early-divergent species *P. parva*. The stem age of the species (*ca.* 4.02 Ma) coincides with a period of rapid woodland expansion, which may have driven this unidirectional xeric adaptation (e.g., Hugall et al., 2008; Rix et al., 2021). *Panesthia parva* displays a unique ecology among the genus, residing in dry, dead treetops and subsisting for months without moisture (J.A. Walker, pers. obs.). Potentially, the associated physiological and behavioral adaptations prohibit a reversion to inhabiting rainforest.

Our results suggest that most species arose through vicariance as mesic forests retracted. The two most speciose clades were found to occur primarily in the Northern and Central Queensland bioregions (Clades B+C; Figure 3). The boundary between these two regions corresponds to the Saint Lawrence Gap, an expanse of dry grassland that forms an arid barrier to dispersal in mesic organisms (Bryant and Krosch, 2016). While the timing of its formation is not resolved with high precision, a number of phylogeographic studies have detected divergences across the gap that date to the mid Pliocene (Baker et al., 2008; Burke et al., 2013; Chapple et al., 2011). This timeframe corresponds to the divergence between the two *Panesthia* clades (*ca.* 3.87 Ma) and suggests that they may have been isolated by the grasslands’ formation. Ecological surveys undertaken by the authors have failed to detect *Panesthia* within the gap, indicating that it remains a barrier to this day.

Within the two clades, OTUs were found to typically occupy small habitat ranges, spanning one or few patches of rainforest or fringing open forest (i.e., *P. ancaudellioides*, *P.* sp. Cape Upstart, *P.* sp. Brandy Creek, *P. cribrata* North, *P. matthewsi*; Figure 2). Micro-endemism is common in dispersal-limited invertebrates, which can persist long-term in small forest isolates (e.g., Harvey et al., 2017; Oberski et al., 2018). Based on our time-calibrated phylogeny, most speciation occurred during the Pliocene (*ca.* 2.5–5 Ma). During this period, intense glacial and interglacial cycles are thought to have caused substantial and rapid retractions in rainforest (Hill, 1994; White, 1986, 1994), producing the highly fragmented distribution seen today. In concordance, species-level structure dated to the Pliocene has been observed in many mesic lineages across the eastern seaboard (Baker et al., 2008; Heimburger et al., 2022; Lucky, 2011; Moreau et al., 2015; Mutton et al., 2019; Ponniah and Hughes, 2004; Sota et al., 2005).

In contrast to the fine-scale endemism observed in northern and central Queensland, the southern reaches of the group’s range are occupied by *P. cribrata* Central + South, which span larger, contiguous distributions from southern Queensland to Victoria. The boundary between the two corresponds to a region of dry sclerophyll, which presumably limits secondary contact (the Brisbane Valley Barrier; Bryant and Krosch, 2016). While present sampling did not cover the complete range of *P. cribrata* South, genetic relationships within the OTUs were not consistently arranged by geographic locality or habitat type, and close relationships were found between samples from distant rainforest and dry forest sites (e.g., Mogo, Capertee Valley and Kempsey). This contrasts with the stricter geographic clustering and deeper structure seen in co-occurring rainforest invertebrates (Garrick et al., 2004; Garrick et al., 2008; Garrick et al., 2012; Symula et al., 2008) and suggests ongoing gene flow between wet and dry sclerophyll populations. Thus, our findings suggest that both niche-conserved vicariance and more recent dispersal through drier forest have contributed to the contemporary distribution of the *Panesthia*.

Finally, secondary calibration with date estimates from Beasley-Hall et al. (2021b) yielded a substantially deeper evolutionary timescale. Under this scenario, the speciation of the *Panesthia* would have occurred against a backdrop of more gradual Miocene aridification, associated with the incipient fragmentation of east-coast rainforests (Byrne et al., 2011). A number of studies have estimated comparable (e.g., Lucky, 2011; Rix and Harvey, 2012; Tallowin et al., 2019) – or older (Gunter et al., 2019; Oberski et al., 2018) – species ages across the eastern seaboard, indicating that some or all of the biogeographic barriers between *Panesthia* clades may have formed during the Miocene. Thus, we cannot rule out this alternative explanation for the group’s evolution.

### 4.3. Colonization of the Lord Howe Island Group

The Lord Howe Island cockroach *P. lata* is one of the most poorly understood members of the genus. Previous studies have only included one (BeasleyLHall et al., 2021b; Legendre et al., 2017; Legendre et al., 2015) or two (Lo et al., 2016) representatives of the species, leaving open questions regarding the timing, route and number of dispersals to the LHIG. By sampling across the full spatial extent of the LHIG, we robustly resolve *P. lata* as a monophyletic group and evidence the species reaching the islands in a single dispersal event. Our analyses also reveal appreciable population structure across the archipelago, which we discuss elsewhere in a focused population genetic investigation of the species (Adams et al., in prep.).

The LHIG is highly remote, and the ancestors of *P. lata* presumably arrived by rafting. Range reconstructions based on the mitogenomic topology were unable to clearly estimate a biogeographic region of origin; however, the species was united with *P. cribrata* South, which is found in New South Wales and far-southern Queensland. The LHIG occurs due east of Port Macquarie, New South Wales, thus a parsimonious explanation is that the islands were colonized from a nearby section of the mainland. However, we note that the phylogenies estimated from nuclear or concatenated mito-nuclear markers instead unite *P. lata* with species occurring further north in Queensland. In which case, propagules could potentially have been transported southwards by surface currents, following the establishment of the East Australian Current at least by the mid-Pliocene (Christensen et al., 2021; Przeslawski et al., 2011). Although biogeographic studies of LHIG terrestrial fauna are scarce, a sister relationship with Queensland species has been found in endemic *Armadillo* isopods (Lillemets and Wilson, 2002) and peloridiid moss bugs (Burckhardt, 2009). Similar patterns are also observed across a large range of marine taxa (e.g., Colgan and Woods, 2022; Veron and Done, 1979; Williams et al., 2011a).

*Panesthia lata* is notable for its tolerance of the uniquely exposed conditions on the small islets surrounding Lord Howe Island proper. Yet, even when accounting for its discordant placement between topologies, *P. lata* was consistently nested among rainforest or sclerophyllous lineages, and its ancestral habitat was inferred to be either closed or open forest. This suggests that the species initially established in rainforest, which is widespread on the main island, before subsequently expanding its environmental tolerance. Due to the non-monophyly of island populations within *P. lata* (Figure 1), it is challenging to discern precisely when and how the niche shift(s) occurred. Potentially, the species may have expanded its environmental tolerance early in its evolutionary history, and subsequently spread to the drier, more exposed islets. The label accompanying the samples from Ball’s Pyramid, which diverged early in the species’ history, indicates that they were collected from leaf litter (an unusually dry habitat for *Panesthia*). Alternatively, it is also plausible that niche expansions occurred multiple times in parallel, in association with the isolation of each islet lineage. Further study is underway to investigate ecological differences among these populations and to refine hypotheses regarding their evolution.

### 4.4. Wing morphology

The *Panesthia* display a broad range of wing morphologies, spanning macropterous, brachypterous and micropterous forms. Even when accommodating the possibility of wing re-evolution (Forni et al., 2022), our reconstructions estimate that wing reduction has occurred 6–8 times independently, including at least twice within *P. cribrata* Central. Given the recency of diversification, possible explanations for the frequent wing reduction include *de novo* mutation, selection on standing polymorphism, or phenotypic plasticity. However, inconsistently with plasticity, wing morphs were strictly partitioned between allopatric populations, with no polymorphism at any single sampling locality. Long-term culture of *Panesthia* has also found wing morphology to be stable across generations (>10 years; H.A. Rose, J.A. Walker, pers. obs.). Therefore, it is likely that each instance of flight loss is independent, corroborating previous reports of frequent wing loss across the subfamily Panesthiinae (Bell et al., 2007b; Roth, 1977, 1979a, b, 1982).

The tendency towards wing reduction is presumably tied to the saproxylic niche. Due to the energetic maintenance costs of flight apparatus, wings are strongly selected against and frequently lost in confined habitats such as logs (Bell et al., 2007b; Roff, 1990). Likewise, the manual shedding of wings in macropterous lineages is ubiquitous, and presumably alleviates energetic costs through the histolysis of flight muscles (Roff, 1989; Tanaka, 1994). Hence, it is unclear how fitness varies between facultatively flightless (macropterous, wing-shed) and permanently flightless (wing-reduced) morphs, and whether any environmental pressures encourage permanent wing reduction.

One potential correlate that unites wing-reduced lineages is their occupation of small habitat patches. In addition to the island-endemic *P. lata*, all mainland flightless lineages have disjunct ranges spanning one or few mesic isolates. A long-standing hypothesis suggests that flight is selected against in insular habitats, to reduce dispersal into surrounding, unsuitable environments (reviewed by Waters et al., 2020). However, it is unknown how frequently, or how far, macropterous cockroaches fly prior to shedding. Measuring dispersal ability prior to wing-shedding would be needed to clarify the potentially deleterious nature of this trait in insular habitats. Likewise, habitat insularity may co-vary with other potentially relevant variables such as temperature, altitude or environmental stability (Roff, 1990, 1994), which were not considered in our habitat reconstructions. The presence of wing polymorphism in *P. cribrata* Central provides an exemplary model system in which to compare closely related, morphologically divergent populations. Fitness assays of different wing morphs of *P. cribrata* Central, complemented by more granular ecological and genomic analyses, could illuminate the underpinnings of this striking parallel evolution.

### 4.5. Conclusions

This study provides new insights into the diversity of the *Panesthia*, revealing 11 divergent genetic lineages. Niche reconstructions suggest that the ancestors of the group occupied closed mesic forest and subsequently speciated through both vicariance and transitions into drier forests as rainforests retracted during the Pliocene. The retraction of rainforest into insular refugia may have also driven parallel wing reduction by exerting selective pressure against flight, although further work is required to test this hypothesis. Our findings further reveal that *P. lata* most likely reached the LHIG in a single colonization event, and that the species expanded into a drier niche after establishment on the archipelago. Overall, our integration of mainland and island taxa offers a holistic view of historical biogeography across two disparate geographic realms.

## Acknowledgements

This project was supported by an Australia Pacific Science Foundation grant (APSF22029), and samples were collected under permits from NSW National Parks and Wildlife Services (SL102663) and the Lord Howe Island Board (05/22). The authors acknowledge the Sydney Informatics Hub and the use of the University of Sydney’s high performance computing cluster, Artemis, for computational resources that contributed to results in the present paper. We are grateful to the Lord Howe Island Board for support and assistance with fieldwork, and to Chris Reid, Derek Smith, Jude Philip and Matthew Hahn for access to historical specimens. Finally, we thank Toby Kovacs for valuable comments on an earlier draft of the manuscript.

## Data availability

The genomic sequences generated in this study are to be uploaded to GenBank prior to publication, and accession numbers will be provided in Supplementary Table S1.

