## Supplementary material for "Plio-Pleistocene decline of mesic forest underpins diversification in a clade of Australian *Panesthia* cockroaches"

**Materials and methods**

**Supplementary Table S1.** List of ingroup and outgroup taxa included in the study, with corresponding Genbank accession numbers. Dashes (-) indicate missing sequences or geographic data, asterisks (*) indicate incomplete mitochondrial genomes for which gene-specific accession numbers are provided in Supplementary Table S2. Australian region abbreviations: Lord Howe Island Group (LHIG), New South Wales (NSW), Queensland (QLD).

| **Taxon** | **Locality / source** | **MA #** | **Sample ID** | **Latitude (° E)** | **Longitude (° S)** | **Mitochondrial genome** | **Nuclear ribosomal operon** |
| --- | --- | --- | --- | --- | --- | --- | --- |
| *Panesthia ancaudellioides* | Annan River, QLD | 163 | JAW-050 | ~15.82 | ~145.25 | TBA | TBA |
| *Panesthia ancaudellioides* | Annan River, QLD | 164 | JAW-051 | ~15.82 | ~145.25 | TBA | TBA |
| *Panesthia ancaudellioides* | Homerule, QLD | 053 | HAR-019 | 15.755 | 145.290 | TBA | TBA |
| *Panesthia ancaudellioides* | Homerule, QLD | 052 | HAR-019b | 15.755 | 145.290 | TBA | TBA |
| *Panesthia ancaudellioides* | Mount Windsor Tablelands, QLD | 165 | JAW-053 | ~16.26 | ~145.05 | TBA | TBA |
| *Panesthia ancaudellioides* | Mount Windsor Tablelands, QLD | 166 | JAW-052 | ~16.26 | ~145.05 | TBA | TBA |
| *Panesthia ancaudellioides* | Unknown | 170 | JAW-057 | N/A | | TBA | TBA |
| *Panesthia ancaudellioides* | Unknown | 171 | JAW-058 | N/A | | TBA | TBA |
| *Panesthia ancaudellioides* | Unknown | 187 | RB4,R3,9 | N/A | | TBA | TBA |
| *Panesthia cribrata* Central | Amamoor, QLD | 161 | HAR-010b | 26.361 | 152.634 | TBA | TBA |
| *Panesthia cribrata* Central | Amamoor, QLD | 162 | HAR-010c | 26.361 | 152.634 | TBA | TBA |
| *Panesthia cribrata* Central | Cameron's Scrub, QLD | 167 | JAW-054 | ~27.49 | ~152.73 | TBA | TBA |
| *Panesthia cribrata* Central | Cooran, QLD | 051 | HAR-014 | 26.262 | 152.830 | TBA | TBA |
| *Panesthia cribrata* Central | Cooran, QLD | 050 | HAR-014b | 26.262 | 152.830 | TBA | TBA |
| *Panesthia cribrata* Central | Gayndah, QLD | 101 | RB1,R3,10 | 25.420 | 151.260 | TBA | TBA |
| *Panesthia cribrata* Central | Gayndah, QLD | 102 | B2,25b | 25.703 | 151.426 | TBA | TBA |
| *Panesthia cribrata* Central | Gayndah, QLD | 103 | B2,25c | 25.703 | 151.426 | TBA | TBA |
| *Panesthia cribrata* Central | Imbil, QLD | 116 | HAR-015 | ~ 24.46 | ~152.68 | TBA | TBA |
| *Panesthia cribrata* Central | Kalpowar, QLD | 091 | HAR-012b | 24.685 | 151.340 | TBA | TBA |
| *Panesthia cribrata* Central | Kalpowar, QLD | 092 | HAR-012c | 24.685 | 151.340 | TBA | TBA |
| *Panesthia cribrata* Central | Kingaroy, QLD | 084 | RB4,R1,4b | 26.703 | 151.779 | TBA | TBA |
| *Panesthia cribrata* Central | Kingaroy, QLD | 085 | RB4,R1,4c | 26.703 | 151.779 | TBA | TBA |
| *Panesthia cribrata* Central | Kroombit Tops, QLD | 123 | HAR-018 | 24.397 | 151.045 | TBA | TBA |
| *Panesthia cribrata* Central | Kroombit Tops, QLD | 121 | HAR-018b | 24.397 | 151.045 | TBA | TBA |
| *Panesthia cribrata* Central | Kroombit Tops, QLD | 122 | HAR-018c | 24.397 | 151.045 | TBA | TBA |
| *Panesthia cribrata* Central | Kroombit Tops, QLD | 109 | B2,22b | 24.383 | 151.003 | TBA | TBA |
| *Panesthia cribrata* Central | Kroombit Tops, QLD | 110 | RB1,R3,8b | 24.370 | 150.940 | TBA | TBA |
| *Panesthia cribrata* Central | Lake Borumba, QLD | 066 | HAR-013b | 26.529 | 152.569 | TBA | TBA |
| *Panesthia cribrata* Central | Lake Borumba, QLD | 067 | HAR-013c | 26.529 | 152.569 | TBA | TBA |
| *Panesthia cribrata* Central | Mapleton, QLD | 149 | HAR-020 | 26.625 | 152.843 | TBA | TBA |
| *Panesthia cribrata* Central | Mapleton, QLD | 147 | HAR-020b | 26.625 | 152.843 | TBA | TBA |
| *Panesthia cribrata* Central | Mount Glorious, QLD | 044 | RB1,R4,20 | N/A | | TBA | TBA |
| *Panesthia cribrata* Central | Mount Glorious, QLD | 034 | HAR-006 | 27.294 | 152.750 | TBA | TBA |
| *Panesthia cribrata* Central | Murgon, QLD | 094 | B2,17b | 26.155 | 151.911 | TBA | TBA |
| *Panesthia cribrata* Central | Murgon, QLD | 095 | B2,17c | 26.155 | 151.911 | TBA | TBA |
| *Panesthia cribrata* Central | Noosa, QLD | 045 | RB1,R3,4 | ~26.38 | ~153.10 | TBA | TBA |
| *Panesthia cribrata* Central | Redwood Park, QLD | 087 | HAR-005 | 27.564 | 151.998 | TBA | TBA |
| *Panesthia cribrata* Central | Redwood Park, QLD | 086 | HAR-005b | 27.564 | 151.998 | TBA | TBA |
| *Panesthia cribrata* Central | Yarraman, QLD | 130 | RB4,R1,2 | ~26.84 | ~151.98 | TBA | TBA |
| *Panesthia cribrata* Central | Yarraman, QLD | 127 | RB4,R1,2b | ~26.84 | ~151.98 | TBA | TBA |
| *Panesthia cribrata* Central | Yarraman, QLD | 100 | HAR-021 | 24.527 | 151.471 | TBA | TBA |
| *Panesthia cribrata* North | Bulburin, QLD | 099 | HAR-021b | 24.527 | 151.471 | TBA | TBA |
| *Panesthia cribrata* North | Bulburin, QLD | 143 | HAR-021 | 24.518 | 151.463 | TBA | TBA |
| *Panesthia cribrata* North | Bulburin, QLD | 142 | HAR-021b | 24.518 | 151.463 | TBA | TBA |
| *Panesthia cribrata* North | Unknown | 033 | HAR-011 | N/A | | TBA | TBA |
| *Panesthia cribrata* North | Goodnight Scrub, QLD | 111 | B2,23b | 25.284 | 151.924 | TBA | TBA |
| *Panesthia cribrata* North | Goodnight Scrub, QLD | 112 | B2,23c | 25.284 | 151.924 | TBA | TBA |
| *Panesthia cribrata* North | Goodnight Scrub, QLD | 113 | B2,23d | 25.284 | 151.924 | TBA | TBA |
| *Panesthia cribrata* North | Goodnight Scrub, QLD | 114 | RB4,R2,20b | 25.228 | 151.922 | TBA | TBA |
| *Panesthia cribrata* North | Goodnight Scrub, QLD | 115 | RB4,R2,20c | 25.228 | 151.922 | TBA | TBA |
| *Panesthia cribrata* North | Gurgeena, QLD | 108 | HAR-017 | 25.451 | 151.383 | TBA | TBA |
| *Panesthia cribrata* North | Gurgeena, QLD | 124 | RB4,R2,18b | 25.453 | 151.385 | TBA | TBA |
| *Panesthia cribrata* North | Many Peaks, QLD | 137 | RB4,R2,19b | 24.527 | 151.474 | TBA | TBA |
| *Panesthia cribrata* North | Monto, QLD | 117 | B2,29b | 24.899 | 151.023 | TBA | TBA |
| *Panesthia cribrata* North | Monto, QLD | 118 | B2,29c | 24.899 | 151.023 | TBA | TBA |
| *Panesthia cribrata* North | Monto, QLD | 119 | B2,30b | 24.899 | 151.342 | TBA | TBA |
| *Panesthia cribrata* North | Monto, QLD | 120 | B2,30c | 24.899 | 151.342 | TBA | TBA |
| *Panesthia cribrata* North | Monto, QLD | 096 | HAR-016b | 24.893 | 151.333 | TBA | TBA |
| *Panesthia cribrata* North | Monto, QLD | 097 | HAR-016c | 24.893 | 151.333 | TBA | TBA |
| *Panesthia cribrata* South | Capertee Valley, NSW | 177 | JAW-064 | N/A | | TBA | TBA |
| *Panesthia cribrata* South | Capertee Valley, NSW | 178 | JAW-065 | N/A | | TBA | TBA |
| *Panesthia cribrata* South | Cheltenham, NSW | 035 | RB1,R4,5 | ~33.76 | ~151.07 | TBA | TBA |
| *Panesthia cribrata* South | Cobbitty, NSW | 046 | RB1,R1,10 | ~34.02 | ~150.67 | TBA | TBA |
| *Panesthia cribrata* South | Cobbitty, NSW | 047 | RB1,R4,6 | ~34.02 | ~150.67 | TBA | TBA |
| *Panesthia cribrata* South | Dorrigo, NSW | 058 | B2,18b | 30.373 | 152.725 | TBA | TBA |
| *Panesthia cribrata* South | Dorrigo, NSW | 059 | B2,18c | 30.373 | 152.725 | TBA | TBA |
| *Panesthia cribrata* South | Ebor, NSW | 060 | RB4,R1,12 | ~30.40 | ~152.34 | TBA | TBA |
| *Panesthia cribrata* South | Ebor, NSW | 061 | B2,19b | ~30.40 | ~152.34 | TBA | TBA |
| *Panesthia cribrata* South | Gosford, NSW | 179 | RB1,R4,7b | 33.440 | 151.313 | TBA | TBA |
| *Panesthia cribrata* South | Gosford, NSW | 180 | RB1,R4,7c | 33.440 | 151.313 | TBA | TBA |
| *Panesthia cribrata* South | Kangaroo Valley, NSW | 055 | RB4,R1,11 | 34.741 | 150.539 | TBA | TBA |
| *Panesthia cribrata* South | Kangaroo Valley, NSW | 054 | RB4,R1,11b | 34.741 | 150.539 | TBA | TBA |
| *Panesthia cribrata* South | Kempsey, NSW | 042 | RB1,R4,9 | 31.533 | 152.788 | TBA | TBA |
| *Panesthia cribrata* South | Killara, NSW | 062 | RB1,R4,3 | 33.768 | 151.145 | TBA | TBA |
| *Panesthia cribrata* South | Killara, NSW | 063 | RB4,R1,11b | 33.768 | 151.145 | TBA | TBA |
| *Panesthia cribrata* South | Killarney, NSW | 138 | B2,21b | 28.279 | 152.446 | TBA | TBA |
| *Panesthia cribrata* South | Killarney, NSW | 139 | RB1,R3,6b | 28.286 | 152.324 | TBA | TBA |
| *Panesthia cribrata* South | Kowmung River, NSW | 048 | RB4,R1,4b | ~34.04 | ~150.19 | TBA | TBA |
| *Panesthia cribrata* South | Kowmung River, NSW | 049 | RB4,R1,4c | ~34.04 | ~150.19 | TBA | TBA |
| *Panesthia cribrata* South | Meadowbank, NSW | 037 | RB1,R4,1 | 33.821 | 151.090 | TBA | TBA |
| *Panesthia cribrata* South | Megalong Valley, NSW | 140 | MV_crib1 | N/A | | TBA | TBA |
| *Panesthia cribrata* South | Megalong Valley, NSW | 141 | MV_crib2 | N/A | | TBA | TBA |
| *Panesthia cribrata* South | Mogo, NSW | 064 | RB4,R1,6 | 35.768 | 150.064 | TBA | TBA |
| *Panesthia cribrata* South | Mogo, NSW | 065 | RB1,R4,15b | 35.768 | 150.064 | TBA | TBA |
| *Panesthia cribrata* South | Mount Coricudgy, NSW | 175 | JAW-062 | ~32.82 | ~150.35 | TBA | TBA |
| *Panesthia cribrata* South | Mount Coricudgy, NSW | 176 | JAW-063 | ~32.82 | ~150.35 | TBA | TBA |
| *Panesthia cribrata* South | Norfolk Island | 074 | HAR-003b | 29.014 | 167.951 | TBA | TBA |
| *Panesthia cribrata* South | Norfolk Island | 075 | HAR-003c | 29.014 | 167.951 | TBA | TBA |
| *Panesthia cribrata* South | Norfolk Island | 088 | RB4,24b | 29.026 | 167.940 | TBA | TBA |
| *Panesthia cribrata* South | Norfolk Island | 089 | RB4,24c | 29.026 | 167.940 | TBA | TBA |
| *Panesthia cribrata* South | Norfolk Island | 090 | RB4,R1,18b | 29.004 | 167.944 | TBA | TBA |
| *Panesthia cribrata* South | Oatley, NSW | 077 | RB1,R4,4c | ~33.98 | ~151.08 | TBA | TBA |
| *Panesthia cribrata* South | Olinda, NSW | 174 | JAW-061 | ~32.82 | ~150.15 | TBA | TBA |
| *Panesthia cribrata* South | Ryde, NSW | 040 | RB3,R3,9 | 33.810 | 151.104 | TBA | TBA |
| *Panesthia cribrata* South | Ryde, NSW | 041 | RB1,R4,2 | 33.810 | 151.104 | TBA | TBA |
| *Panesthia cribrata* South | Tallebudgera Creek, QLD | 031 | RB1,R3,1 | 28.228 | 153.313 | TBA | TBA |
| *Panesthia cribrata* South | Tallebudgera Creek, QLD | 032 | RB4,R3,1 | 28.228 | 153.313 | TBA | TBA |
| *Panesthia cribrata* South | Taree, NSW | 079 | B2,16c | ~31.98 | ~152.52 | TBA | TBA |
| *Panesthia lata* | Ball's Pyramid, LHIG | 001 | K.487941 | N/A | | TBA | TBA |
| *Panesthia lata* | Ball's Pyramid, LHIG | 002 | K.487942 | N/A | | TBA | TBA |
| *Panesthia lata* | Blackburn Island, LHIG | 151 | PL3 | 31.536 | 159.072 | TBA | TBA |
| *Panesthia lata* | Blackburn Island, LHIG | 152 | PL4 | 31.536 | 159.072 | TBA | TBA |
| *Panesthia lata* | Blackburn Island, LHIG | 153 | PL6 | 31.536 | 159.072 | TBA | TBA |
| *Panesthia lata* | Blackburn Island, LHIG | 154 | PL7 | 31.536 | 159.072 | TBA | TBA |
| *Panesthia lata* | Blackburn Island, LHIG | 155 | PL17 | 31.536 | 159.072 | TBA | TBA |
| *Panesthia lata* | Lord Howe Island, LHIG | 008 | K.487922 | N/A | | TBA | TBA |
| *Panesthia lata* | Lord Howe Island, LHIG | 018 | K.487932 | N/A | | TBA | TBA |
| *Panesthia lata* | Lord Howe Island, LHIG | 188 | NHEN.66552 | N/A | | TBA | TBA |
| *Panesthia lata* | Lord Howe Island, LHIG | 189 | NHEN.66559 | N/A | | TBA | TBA |
| *Panesthia lata* | Lord Howe Island, LHIG | 190 | NHEN.66554 | N/A | | TBA | TBA |
| *Panesthia lata* | Lord Howe Island, LHIG | 191 | NHEN.66555 | N/A | | TBA | TBA |
| *Panesthia lata* | North Bay, LHIG | 157 | PL2 | 31.517 | 159.042 | TBA | TBA |
| *Panesthia lata* | North Bay, LHIG | 159 | PL18 | 31.517 | 159.042 | TBA | TBA |
| *Panesthia lata* | North Bay, LHIG | 160 | PL19 | 31.517 | 159.042 | TBA | TBA |
| *Panesthia lata* | Roach Island, LHIG | 024 | K.383796 | 31.500 | 159.068 | TBA | TBA |
| *Panesthia lata* | Roach Island, LHIG | 026 | K.383798 | 31.500 | 159.068 | TBA | TBA |
| *Panesthia lata* | Roach Island, LHIG | 027 | K.383799 | 31.502 | 151.069 | TBA | TBA |
| *Panesthia lata* | Roach Island, LHIG | 028 | K.383800 | 31.500 | 159.068 | TBA | TBA |
| *Panesthia lata* | Roach Island, LHIG | 023 | K.487940 | N/A | | TBA | TBA |
| *Panesthia lata* | Roach Island, LHIG | 025 | K.383797 | 31.502 | TBA | TBA | TBA |
| *Panesthia lata* | Roach Island, LHIG | 029 | K.383801 | 31.502 | 151.069 | TBA | TBA |
| *Panesthia matthewsi* | Biggenden, QLD | 150 | RB4,R2,8 | 25.570 | 152.053 | TBA | TBA |
| *Panesthia matthewsi* | Coalstoun Lakes, QLD | 057 | HAR-002 | 25.615 | 151.895 | TBA | TBA |
| *Panesthia matthewsi* | Coalstoun Lakes, QLD | 056 | HAR-002b | 25.615 | 151.895 | TBA | TBA |
| *Panesthia matthewsi* | Mount Walsh, QLD | 144 | B2,51b | 25.570 | 152.053 | TBA | TBA |
| *Panesthia matthewsi* | Mount Walsh, QLD | 145 | B2,51c | 25.570 | 152.053 | TBA | TBA |
| *Panesthia matthewsi* | Mount Walsh, QLD | 070 | HAR-008b | 25.571 | 152.053 | TBA | TBA |
| *Panesthia matthewsi* | Mount Walsh, QLD | 071 | HAR-008c | 25.571 | 152.053 | TBA | TBA |
| *Panesthia parva* | Mareeba, QLD | 172 | JAW-059 | 17.046 | 145.526 | TBA | TBA |
| *Panesthia parva* | Beasley-Hall et al. (2021) |  |  | N/A | |  | - |
| *Panesthia* sp. Airlie Beach | Brandy Creek, QLD | 168 | JAW-055 | 20.341 | TBA | TBA | TBA |
| *Panesthia* sp. Airlie Beach | Brandy Creek, QLD | 169 | JAW-056 | 20.341 | 148.682 | TBA | TBA |
| *Panesthia* sp. Airlie Beach | Brandy Creek, QLD | 184 | B2,50 | 20.341 | 148.678 | TBA | TBA |
| *Panesthia* sp. Airlie Beach | Patullo Road, QLD | 183 | B2,48 | 20.271 | 148.582 | TBA | TBA |
| *Panesthia* sp. Cape Upstart | Cape Upstart, QLD | 181 | JAW-066 | 19.770 | 147.808 | TBA | TBA |
| *Panesthia* sp. Cape Upstart | Cape Upstart, QLD | 185 | RB1,R2,19 | 19.732 | 147.814 | TBA | TBA |
| *Panesthia* sp. Mt Windsor | Mount Windsor Tablelands, QLD | 182 | JAW-067 | 16.259 | 145.049 | TBA | TBA |
| *Panesthia* sp. Koombooloomba | Koombooloomba State Forest, QLD | 199 | K1 | 17.884 | 145.527 | TBA | TBA |
| *Panesthia angustipennis angustipennis* | Che et al. (2022) |  |  | N/A | | - | MZ049968 |
| *Panesthia angustipennis angustipennis* | Wang et al. (2023) |  |  | N/A | | - | OQ738140 |
| *Panesthia angustipennis angustipennis* | Beasley‐Hall et al. (2021) |  |  | N/A | | MW996590 | - |
| *Panesthia angustipennis cognata* | Wang et al. (2023) |  |  | N/A | | OQ736943 | OQ738138 |
| *Panesthia angustipennis spadica* | Kinjo et al. (2021) |  |  | N/A | | OL685387 | - |
| *Panesthia angustipennis yayeyamensis* | Kinjo et al. (2021) |  |  | N/A | | OL685388 | - |
| *Panesthia tryoni tryoni* | Beasley-Hall et al. (2021) |  |  | N/A | | MW996603 | - |

**Supplementary Table S2.** Primers and polymerase chain reaction (PCR) amplification conditions for mitochondrial *CO1* and *16S*.

| **Marker** | **Direction** | **Sequence (5’–3’)** | **PCR protocol** | **Reference** |
| --- | --- | --- | --- | --- |
| *CO1* | Forward  Reverse | GGTCAACAAATCATAAAGATATTGG  TAAACTTCAGGGTGACCAAAAAATCA | Initial denaturation for 2 min at 94 °C; 30 cycles of denaturation at 94 °C for 30 sec, annealing at 65 °C for 30 sec, and extension at 72 °C for 30 sec; and final extension at 72 °C for 10 min | Vrijenhoek (1994) |
| *16S* | Forward  Reverse | CGCCTGTTTATCAAAAACAT  CCGGTCTGAACTCAGATCACG |  | Simon et al. (1994) |

**Supplementary Table S3.** Best-fitting substitution models for each partition in the mitogenomic and nuclear data sets, as selected by ModelFinder (Kalyaanamoorthy et al., 2017). Abbreviations: protein-coding gene (PCG), ribosomal RNA (rRNA), transfer RNA (tRNA).

| **Dataset** | **Partition** | **Size (bp)** | **Substitution model** |
| --- | --- | --- | --- |
| Mitogenome | PCG codon 1 | 3194 | TIM2 + F + I + G4 |
|  | PCG codon 2 | 3194 | TIM2 + F + R3 |
|  | PCG codon 3 | 3194 | TIM + F + I + R3 |
|  | *CO1* | 1532 | TIM2 + F + I + R3 |
|  | rRNA | 2197 | TIM2 + F + I + R3 |
|  | tRNA | 1499 | TPM2u + F + R4 |
| Nuclear ribosomal operon | *28S* | 2767 | TN + F + I |
|  | *5.8S + 18S* | 2157 | JC + I |
|  | *ITS1 + ITS2* | 481 | HKY + F + G4 |

*Species delimitation*

We utilised a two-step approach to investigate evolutionary relationships within the *Panesthia*. First, species hypotheses were generated *de novo* from the mitogenomic data set using GMYC and ASAP. We implemented GMYC using the single-threshold method in the R package *splits* v.1.0-20 (Ezard et al. 2009), applied to the ultrametric phylogeny estimated in BEAST. ASAP was run through the developers’ web server (<https://bioinfo.mnhn.fr/abi/public/asap/>, accessed March 13, 2024) with default settings and a Kimura (1980) two-parameter model of nucleotide substitution. Both analyses were undertaken using a pruned dataset comprising only ingroup samples.

Second, we validated the output delimitations using Bayesian inference in Bayesian Phylogenetics and Phylogeography v.4.1.4 (BPP; Yang 2015). When provided with gene alignments, a “guide” tree topology and putative species boundaries, the program iteratively collapses these boundaries and compares model adequacy under different delimitation schemes. We implemented BPP using a two-partitioned alignment comprising (1) complete mitogenomes and (2) the nuclear ribosomal operon. The 12-species scheme generated by ASAP (i.e., the more speciose result from the species discovery step) and the concatenated mito-nuclear topology were used as a guide tree.

BPP analysis was undertaken using species delimitation method 1 and algorithm 0 (see Yang and Rannala 2010 for formulations). We followed the recommendations in the package to apply diffuse inverse gamma-distributed priors for mutation-scaled population size (Θ = *IG*(3, 0.05)) and age of the root (𝛕 = *IG*(3, 0.012)). Experimentation with a range of prior settings showed results to be highly consistent. Models were run for 1,000,000 steps, sampling every two steps, with a burn-in of 10%. We performed three independent runs to confirm the stability of parameter estimates, and accepted species with posterior probability > 0.95 across all replicates.

**Results**

**Supplementary Table S4.** Results of a topology test performed in IQTREE2, comparing the unconstrained nuclear phylogeny against a constrained nuclear phylogeny following the mitogenomic branching order. Values measure statistical support relative to the best-supported (unconstrained) topology, positive (+) or negative (-) signs indicate whether the p-value was associated with significantly higher or lower support, respectively. ΔL, difference in log-likelihood; bp-RELL, bootstrap proportion using the RELL method (Kishino et al. 1990); c-ELW, expected Likelihood Weight (Strimmer and Rambaut 2002); p-KH, p-value of the one sided Kishino-Hasegawa test (Kishino and Hasegawa 1989); p-SH p-value of the Shimodaira-Hasegawa test (Shimodaira and Hasegawa 1999); p-AU, p-value of the approximately unbiased test (Shimodaira 2002).

| **Tree** | **ΔL** | **bp-RELL** | **c-ELW** | **p-KH** | **p-SH** | **p-AU** |
| --- | --- | --- | --- | --- | --- | --- |
| Unconstrained | 0 | 1 | 1 | 0.998 | 1 | 1 |
| Constrained | 71.257 | <0.001 | <0.001 | 0.002 (-) | 0.002 (-) | <0.001 (-) |

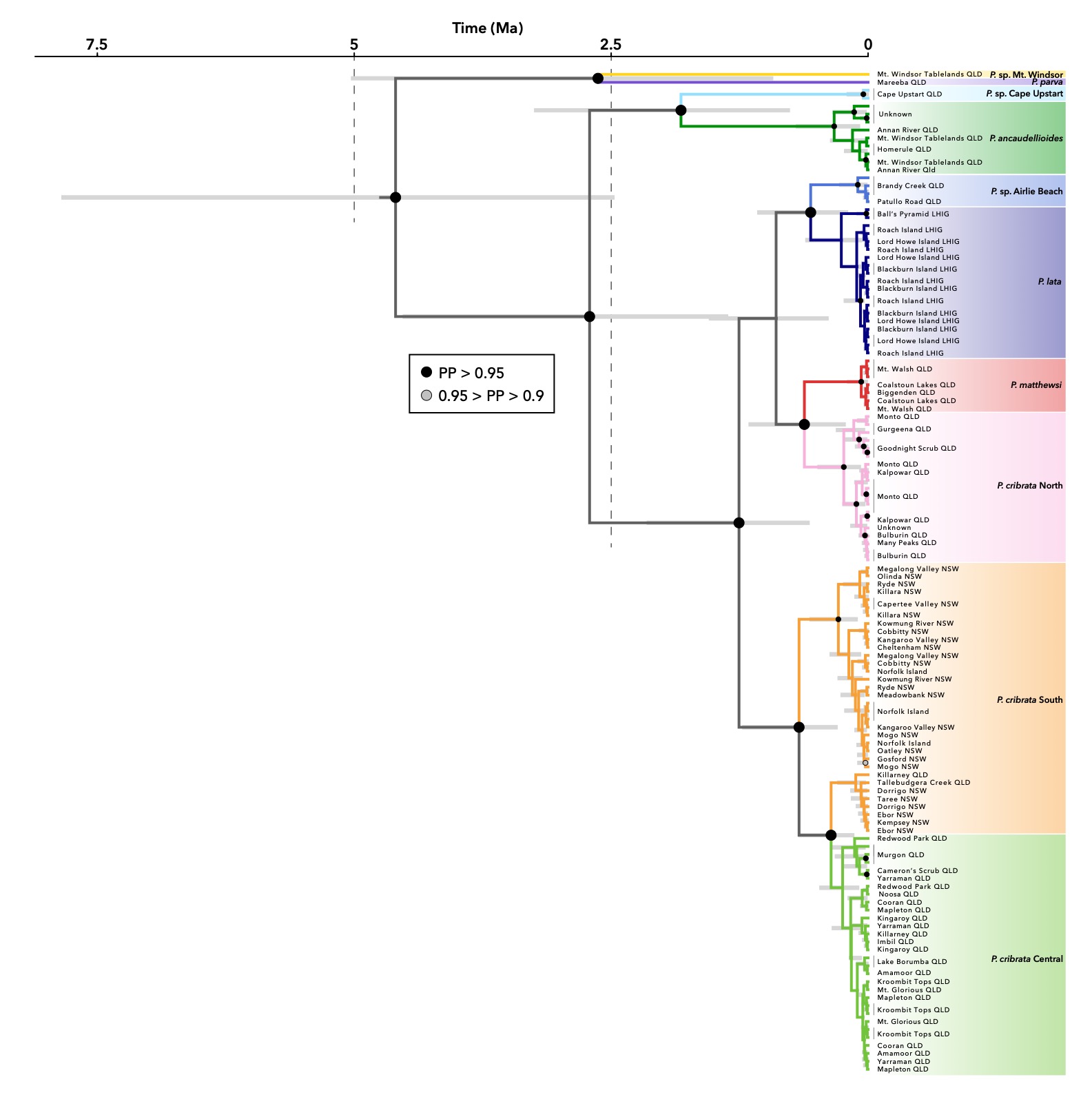

**Supplementary Figure S1.** Dated phylogeny of the *Panesthia* inferred from the nuclear ribosomal operon in BEAST. Only ingroup taxa are shown. The evolutionary timescale was calibrated using a prior estimate for the *28S* rate of evolution (Allegrucci et al., 2011). PP: posterior probability, QLD: Queensland, NSW: New South Wales, LHIG: Lord Howe Island Group. Size of node labels varied for visual clarity. Stars to right of tips denote historical specimens of *Panesthia lata*. Scale axis is in millions of years. Nomenclature and colouration of operational taxonomic units follows the mitogenomic (BPP) results.

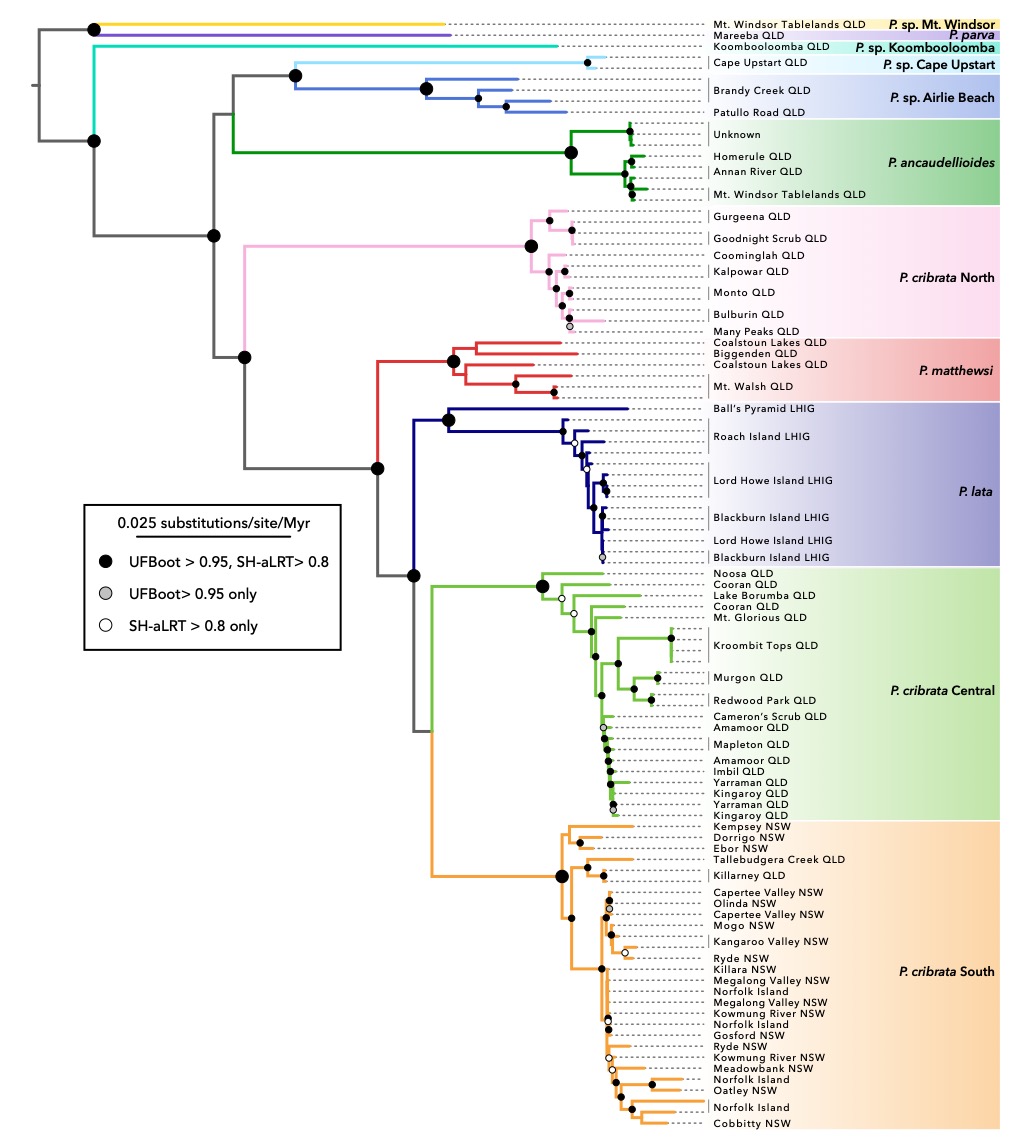

**Supplementary Figure S2.** Maximum-likelihood phylogeny of the *Panesthia* inferred from complete mitochondrial genomes and the nuclear ribosomal operon in IQTREE2. UFBoot: ultrafast bootstrap, SH-aLRT: SH-like likelihood ratio test. Intraspecific nodes not labelled. Nomenclature and colouration of operational taxonomic units follows the mitogenomic (BPP) results.

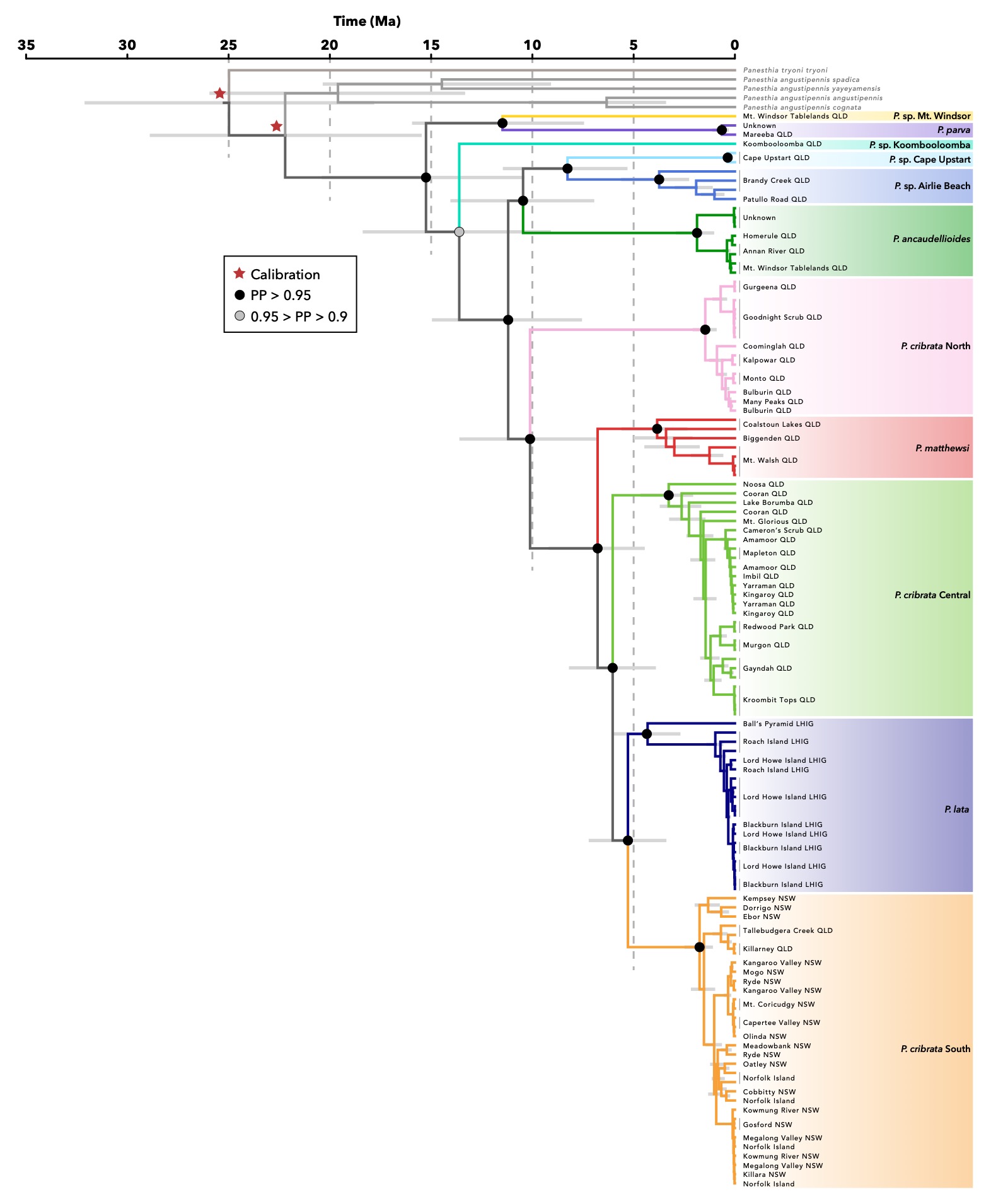

**Supplementary Figure S3.** Dated phylogeny of the *Panesthia* inferred from complete mitochondrial genomes in BEAST. The evolutionary timescale was estimated by applying secondary calibrations to two backbone nodes, based on date estimates in Beasley-Hall et al. (2021b; calibrated nodes indicated by stars). QLD: Queensland, NSW: New South Wales, LHIG: Lord Howe Island Group. Intraspecific nodes not labelled, however support is qualitatively similar to Figure 1. Nomenclature and colouration of operational taxonomic units follows the BPP results.

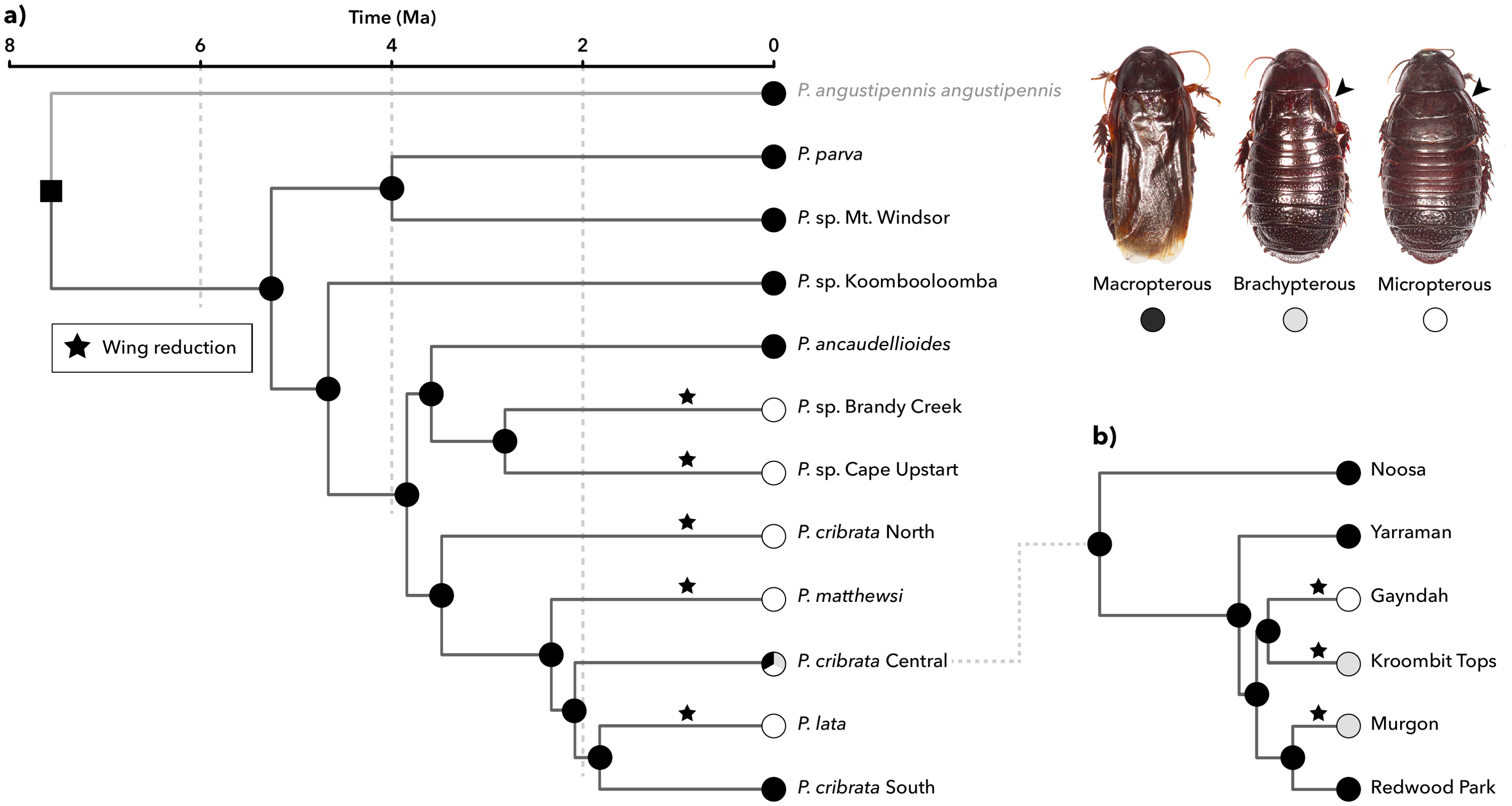

**Supplementary Figure S4.** Evolution of wing morphology with wing re-evolution permitted, reconstructed in *phytools* over the Bayesian chronogram inferred from complete mitogenomes in BEAST. **a)** Ancestral wing morphology of the 11 operational taxonomic units (OTUs). **b)** Ancestral wing morphology within the polymorphic OTU *Panesthia cribrata* Central, pruned to include a representative selection of sampling localities. Circles at nodes indicate the most probable ancestral state and the square at the root node the fixed ancestral state. States at all internal nodes were estimated with probability > 50%. Stars denote inferred wing reduction events. **Inset:** representative habitus images of wing morphs. Arrows indicate reduced forewings. Photographs by Braxton Jones.
